# Development of a genetic evaluation for hair shedding in American Angus cattle to improve thermotolerance

**DOI:** 10.1101/2020.05.21.109553

**Authors:** Harly J. Durbin, Duc Lu, Helen Yampara-Iquise, Stephen P. Miller, Jared E. Decker

**Affiliations:** University of Missouri, Columbia, MO 65211; Angus Genetics Inc., St. Joseph, MO 64506

## Abstract

**Background:** Heat stress and fescue toxicosis caused by ingesting tall fescue infected with the endophytic fungus Epichloë coenophiala represent two of the most prevalent stressors to beef cattle in the United States, costing the beef industry millions of dollars each year. The rate at which a beef cow sheds her winter coat early in the summer is an indicator of adaptation to heat and an economically relevant trait in many parts of the U.S. Further, research suggests that early-summer hair shedding may be reflective of tolerance to fescue toxicosis, as vasoconstriction induced by fescue toxicosis limits the ability of an animal to shed its winter coat. Here, we developed parameters for routine genetic evaluation of hair shedding score in American Angus cattle and identified genomic loci associated with variation in hair shedding score via genome-wide association analysis (GWAA).

**Results:** Hair shedding score was found to be moderately heritable (h2 = 0.34 to 0.40), with differing repeatability estimates between cattle grazing versus not grazing endophyte-infected tall fescue. Our results suggest modestly negative genetic and phenotypic correlations between a dam’s hair shedding score (lower score is earlier shedding) and the weaning weight of her calf, one metric of performance. Together, these results indicate that economic gains can be made via the use of hair shedding score breeding values to select for heat tolerant cattle. GWAA identified 176 variants significant at FDR < 0.05. Functional enrichment analyses using genes within 50 Kb of these variants identified pathways involved in keratin formation, prolactin signaling, host-virus interaction, and other biological processes.

**Conclusions:** This work contributes to a continuing trend in the development of genetic evaluations for environmental adaptation. The results of this work will aid beef cattle producers in selecting more sustainable and climate-adapted cattle, as well as enable the development of similar routine genetic evaluations in other breeds.

## Background

At the beginning of the summer, many mammalian species molt thick winter coats in response to changing day length in order to prepare for warmer temperatures (1–6). There is evidence of quantitative variation in the rate and timing of this yearly shedding across taxa (7,8), including cattle (9). In warm climates, cattle that shed their winter coat earlier and more completely have an adaptive advantage over later-shedding herd-mates. Late-shedding cattle will need to partition energy that could have gone towards growth & production towards overcoming heat stress (10). Economic losses attributable to heat stress cost the U.S. beef cattle industry > $360 million each year in 2003 (11) which equates to ~ $518 million in 2020 after adjustment for inflation. However, there is currently no national-scale genetic evaluation for heat tolerance.

In the United States, much of the beef herd that is at risk of heat stress is also at risk for fescue toxicosis. Tall fescue (*Lolium arundinaceum*) is the most widely available forage in the United States (12), thanks in part to its symbiotic relationship with the endophytic fungus *Epichloë coenophiala*. *Epichloë coenophiala* produces ergot alkaloids that benefit the forage by increasing drought tolerance and pathogen resistance (13), but negatively impact livestock to varying degrees. In cattle, one side-effect of fescue toxicosis is peripheral vasoconstriction, which reduces the animal’s ability to dissipate heat. The ergot alkaloids that cause fescue toxicosis also disrupt the hair follicle growth cycle, which interferes with hair coat shedding and in turn further increases the potential for heat stress (14). Therefore, effective early-summer hair shedding while grazing endophyte-infected (hereafter referred to as “toxic”) tall fescue may also be an indicator of tolerance to fescue toxicosis.

One way to mitigate heat stress is through introgression of beneficial alleles from tropically adapted breeds (15). However, this can take many generations and may come at the cost of other production traits. An alternative strategy is the exploitation of standing genetic variation in the population of interest. Recently, interest has grown in augmenting national genetic evaluations with predictions of regional adaptability and suitability (16–18), particularly through the use of novel traits (19). Here, we develop parameters for a prototype national genetic evaluation of hair shedding in American Angus cattle, a novel trait that directly influences cattle’s ability to dissipate heat. This evaluation will aid beef cattle producers in heat-stressed regions in the selection of more sustainable cattle.

## Methods

### Data

All data originated from purebred cattle registered in the American Angus Association (AAA) and commercial cattle enrolled in the AAA Breed Improvement Record program. Phenotypic data comprised hair shedding scores recorded by beef cattle producers enrolled in the Mizzou Hair Shedding Project (MU data) between 2016 and 2019 in combination with hair shedding scores collected by technicians in 2011, 2012, 2018, and 2019 as part of Angus Foundation-funded projects at Mississippi State University and North Carolina State University (AGI data). Across all years and both datasets, scores were recorded on one day between April 17th and June 30th in the late spring or early summer, with most scores recorded in mid- to late-May. Hair shedding was evaluated using a 1-5 visual appraisal scale, where 1 was 0% dead winter coat remaining and 5 was 100% winter coat remaining based on the systems developed by Turner & Schleger (1960) (20) and Gray et al. (2011) (21) (Figure 1). While there is variation in the onset of hair shedding between individuals, cattle and other mammals tend to shed from the head towards the tail and from the topline towards the legs (2,8,22).

**Figure 1.**
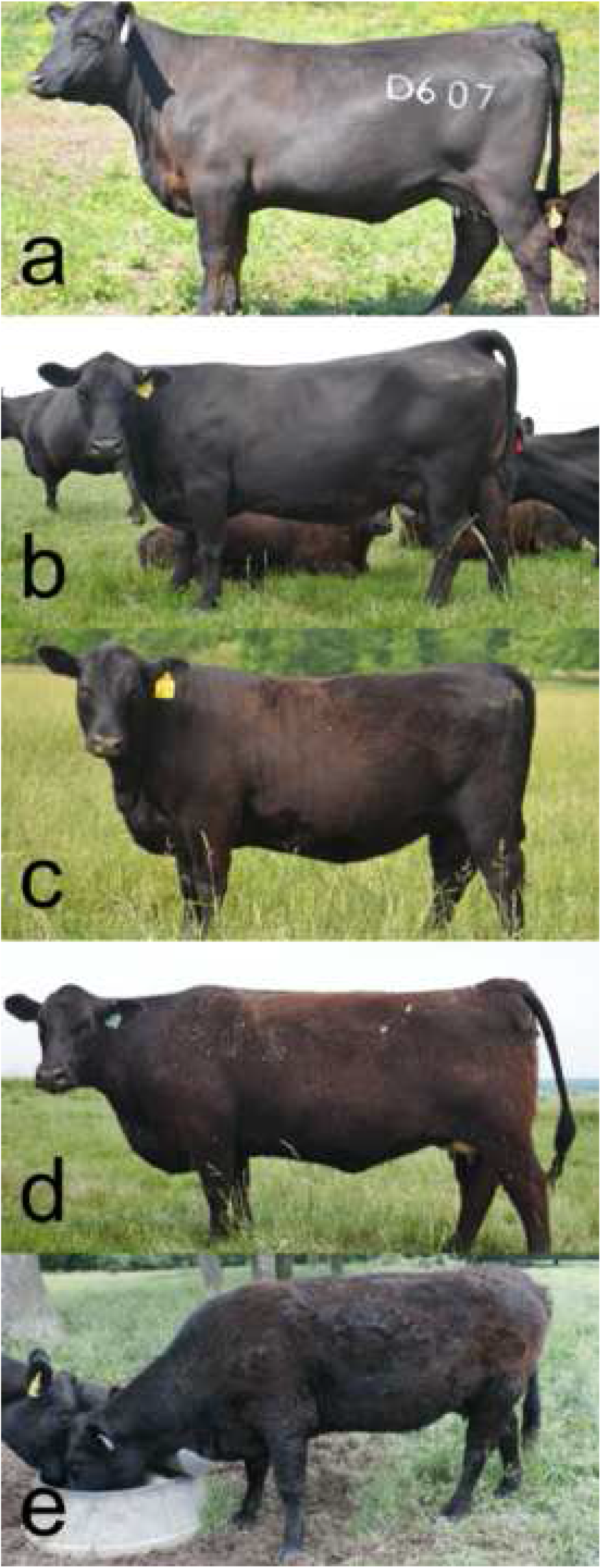
Hair shedding scoring system. Examples of the 1 to 5 visual appraisal hair shedding scoring system used in this research. **a.** Score of 1, 0% dead winter coat remaining. **b.** Score of 2, approximately 25% of winter coat remaining, typically observed on lower hindquarter, flank and belly. **c.** Score of 3, approximately 50% of winter coat remaining. **d.** Score of 4, approximately 75% of winter coat remaining. Hair is typically removed from the head and neck first. **e.** Score of 5, 100% winter coat remaining.

Records were removed when the breeder-reported sex of an animal did not match the sex recorded in the AAA pedigree. Hair shedding scores originating from male animals comprised < 1% of the dataset and only female records were retained. Age classifications were assigned to each record based on age in days determined by the AAA-recorded birth date and the date the hair shedding score was recorded. Similar to the system used in the Beef Improvement Federation (BIF) Guidelines for age-of-dam classification (23), age classifications were defined as *(n*365d) – 90d* to *((n+1)*365d)-90d*, where *n* is the age classification and *d* is days. Records where the breeder-reported age in years differed from the calculated age classification by more than two years and records from animals younger than 275 days-of-age were removed. When no calving season was reported, it was imputed using the most recent natural birth calving date available in the AAA database prior to the recorded score. When no natural birth calving dates were available, calving season was imputed using the animal’s own birth date. In the AGI data, some animals were scored by multiple scoring technicians on the same day. In these cases, phenotypes recorded on the same animal on the same day were averaged. In the MU data, participating producers were asked to report whether or not (yes or no) animals grazed toxic fescue during the spring of the recording year. Grazing status was not explicitly recorded in the AGI data, but animals scored in Texas were assumed not to have grazed toxic fescue. This resulted in 14,839 scores in the combined, filtered dataset. Of 8,619 total individuals, 49% had between 2 and 6 years of data. Most data came from herds in the Southeast and Fescue Belt (Figure 2). The mean hair shedding score was slightly higher in the AGI data (*μ* = 3.10; *n* = 6,374) compared to the MU data (*μ* = 2.65; *n* = 8,465), but the standard deviation was identical in both datasets (*σ* = 1.15).

**Figure 2.**
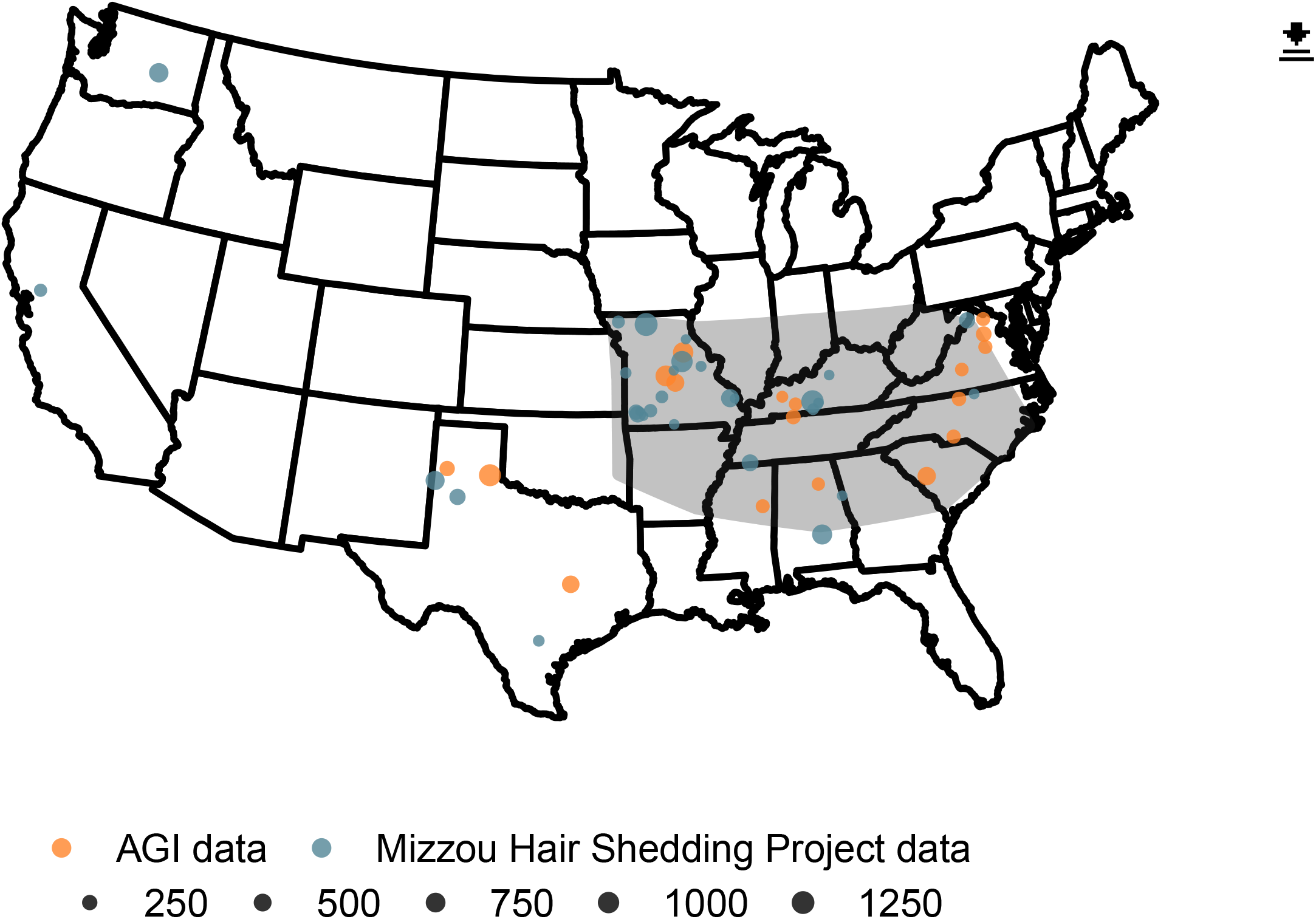
Geographic distribution of animals with hair shedding scores. Hair shedding scores in both the AGI and MU datasets originate primarily from the South and the Fescue Belt. Here, the approximate location of the Fescue Belt is shaded in grey. Size of circles denotes the number of hair shedding scores recorded at that location. Farmers and ranchers in the MU dataset reported whether cattle grazed the predominant endophyte-infected fescue forage or a different forage species.

### Genotypes & imputation

Genotypes for 3,898 animals were imputed to a union marker set of the GGP-F250 genotyping chip and various commercial assays using F-Impute v.3.0 (24). The commercial assays were those used in routine genotyping of Angus cattle for genomic selection purposes, which include ~ 50K markers or a lower density panel that can be imputed to ~ 50K with sufficient accuracy. The GGP-F250 was designed to genotype functional variants and thus has more variants at low minor allele frequencies (25). Therefore, no MAF filter was applied during imputation beyond the removal of monomorphic SNP. Animals and markers with call rates below 85% were removed. The resulting marker set consisted of 174,932 autosomal variants.

### Construction of the blended relationship matrix *H^−1^*

In single-step genomic BLUP as used in the AAA National Cattle Evaluation (NCE), relationships between individuals are represented in the matrix *H^−1^*, a blended form of the genomic and additive relationship matrices (26). In all subsequent models including a random genetic effect, *H^−1^* was constructed using the 3-generation pedigree (17,652 total animals; 1,987 distinct sires and 9,509 distinct dams) in combination with imputed genotypes.

### Contemporary group definition

Appropriate contemporary grouping is fundamental to accurate genetic evaluation, but the power of contemporary grouping is undermined by over-parameterization. In order to determine the best implementation of contemporary grouping in routine genetic evaluation of hair shedding, the effects of several factors were quantified using AIREMLF90 implemented in the BLUPF90 suite of programs (27).

### Age

In order to preliminarily quantify the effect of age on hair shedding score, we fit age-in-years as a fixed effect in a repeated records animal model. Next, two models were fit to determine the best strategy for age classification during contemporary grouping. In the first model, age was classified as 1, 2, 3, or other (hereafter referred to as the “four age classes” model). In the second, age was classified according to the guidelines set by the BIF for age-of-dam effects on birth weight and weaning weight (2, 3, 4, 5-9, 10, 11, 12, 13+; (23)) plus yearlings. Both models were compared using AIC and a likelihood ratio test against a null model with no age effect. In all three models with age in years or age group fit as fixed effects (summarized below), age classifications with fewer than five animals were excluded.

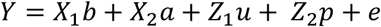

Where:

- *Y* is hair shedding score
- *b* is the effect of contemporary group, defined as farm ID, year, calving season, score group, and toxic fescue grazing status
- *a* is the effect of age group of individual (based on age-in-years, BIF age-of-dam classifications, or four age classes)
- *u* is the additive genetic effect and 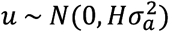
- *p* is the permanent environment effect and 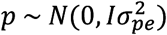
- *e* is the random residual and 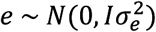
- *X*_1_, *X*_2_, *Z*_1_, and *Z*_2_ are incidence matrices relating the elements of *Y* to *b*, *a*, *u*, and *p*, respectively

### Fescue

Cattle reared in heat-stressed regions but not exposed to endophyte-infected fescue demonstrate similar benefits from early summer hair shedding, but it is unclear if the biological mechanisms governing hair shedding under fescue toxicosis and heat stress alone are the same. This could have implications for routine genetic evaluation, as it might require that some hair shedding score observations be treated as a separate trait. In order to clarify the relationship between hair shedding score while grazing toxic fescue versus while not grazing toxic fescue, we calculated the covariance and genetic correlation between hair shedding score grazing toxic fescue and not grazing toxic fescue using the bivariate repeated records animal model below.

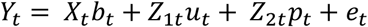

Where:

- *Y* is hair shedding score and *t* is toxic fescue grazing status (yes or no)
- *b* is the effect of contemporary group, defined as farm ID, year, calving season, score group, and age class
- *u* is the additive genetic effect and 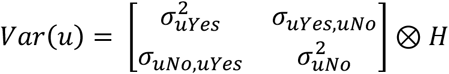
- *p* is the permanent environment effect and 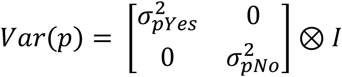
- *e* is the random residual and 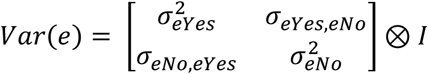
- *X*, *Z*_1_, and *Z*_2_ are incidence matrices relating the elements of *Y* to *b*, *u*, and *p*, respectively Additionally, we tested a univariate model with toxic fescue grazing status fit as a fixed effect.

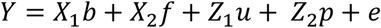

Where:

- *Y* is hair shedding score
- *b* is the contemporary group effect
- *f* is the toxic fescue status effect (yes or no)
- *u* is the additive genetic effect and 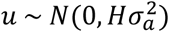
- *p* is the permanent environment effect and 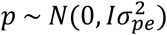
- *e* is the random residual and 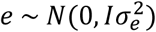
- *X*_1_, *X*_2_, *Z*_1_, and *Z*_2_ are incidence matrices relating the elements of*Y* to *b*, *f*, *u*, and *p*, respectively

In both models, only females with known toxic fescue grazing status were retained for analysis. Contemporary groups were defined as the combination of farm ID, year, calving season, score group, and age class (yearling, 2-year-old, 3-year-old, or other). Contemporary groups with fewer than five animals or no variation were discarded, resulting in 5,832 observations from cattle grazing toxic fescue and 4,197 observations from cattle not grazing toxic fescue. Three hundred ninety-six animals had observations over multiple years both grazing and not grazing toxic fescue.

### Genetic parameters and breeding values

Variance components, heritability, repeatability, and preliminary breeding values were estimated using the repeated records animal model below implemented in AIREMLF90.

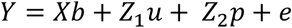

Where:

- *Y* is hair shedding score
- *b* is the contemporary group effect
- *f* is the toxic fescue status effect (yes or no)
- *u* is the additive genetic effect and 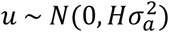
- *p* is the permanent environment effect and 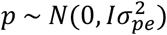
- *e* is the random residual and 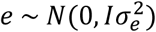
- *X*, *Z*_1_, and *Z*_2_ are incidence matrices relating the elements of *Y* to *b*, *u*, and *p*, respectively

Contemporary groups were defined as a combination of farm, year, calving season (spring or fall) toxic fescue grazing status (yes or no), age group (yearling, 2-year-old, 3-year-old, or other), and score group. In herds where cattle were hair shedding scored over more than one day, the score group was determined using a 7-day sliding window to maximize the number of animals per contemporary group. In the future, it will be recommended that producers hair shedding score all cattle within a week of one another to enable accurate contemporary grouping. Though yearling heifers haven’t yet experienced the stress of pregnancy, calving season/birth season is a good proxy for management group in the absence of breeder-reported codes. Therefore, “calving season” was included in the contemporary group definition for all animals regardless of reproductive status. Contemporary groups with fewer than 5 animals or no variation were dropped. This resulted in 14,438 total scores from 8,449 animals in 395 contemporary groups.

In order to evaluate model bias, we estimated breeding values in ten separate iterations, excluding all phenotypes for a randomly selected 25% of animals. These “partial” breeding values were then compared to breeding values obtained via the “whole” model including all possible information using several parameters proposed by (28). First, we calculated the absolute difference between whole breeding values and partial breeding values for the validation set, or animals whose phenotypes were excluded 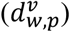 and the reference set, or animals whose phenotypes were not excluded 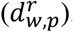. The expectation of this value is zero in the absence of bias, where bias is introduced by incorrect estimation of the genetic trend. Next, we regressed whole breeding values on partial breeding values for both validation 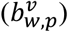 and reference 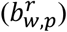 sets. In this model, deviations of the slope from one are suggestive of dispersion. Finally, we calculated the correlation between partial and whole breeding values 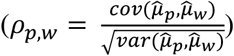 within the validation and reference sets, where the correlation within the validation set 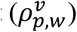 is a metric of prediction accuracy.

### Weaning weight

Weaning weight phenotypes and contemporary group designations came from the weekly growth run of the AAA NCE. Prior to entering the NCE, phenotypes were adjusted for age-of-dam effects as used in Angus’s weekly NCE and to 205 days-of-age. Weaning weight data were aggregated for cows with at least one hair shedding score recorded, all of their recorded calves, their weaning weight contemporary group peers, and all of their recorded calves’ weaning weight contemporary group peers. Weaning weights from calves born via embryo transfer and contemporary groups with fewer than five animals or no variation were excluded, resulting in 40,794 total weaning weight and 14,039 total hair shedding score records. Of the 45,420 phenotyped animals retained for analysis, 3,850 had both a recorded weaning weight and at least one hair shedding score.

Conceivably, environmental factors affecting a dam’s hair shedding performance could also affect the direct weaning weight of her calf and her maternal effect on the calf’s growth, creating a residual covariance between the two traits. In order to reflect this covariance, a bivariate model was fit in which a direct weaning weight effect was modeled for the calf, a maternal weaning weight effect was modeled for the cow, and a direct hair shedding score effect was modeled for the cow. In practice, this model was implemented by fitting a direct and maternal genetic effect for weaning weight, a maternal genetic effect for hair shedding, and no direct genetic effect of hair shedding (no effect of the calf on the hair shedding score of its dam). This model created a direct tie between a dam’s hair shedding score and the calf she weaned that year, which more accurately reflects the relationship of interest and is similar to models used to assess the correlations between weaning weight and actual milk yield (29). For cows with hair shedding scores but no calf weaning weight reported during the scoring year, a “dummy calf” with a weaning weight set to missing and unknown sire was created. This model was fit 3 different times: once including only dams explicitly recorded to have been grazing toxic fescue, once including only dams explicitly recorded to have not been grazing toxic fescue, and once with all available data.

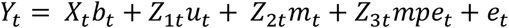

Where:

- *Y*_*t*_ is the phenotype and *t* is the trait (hair shedding score [HS] or weaning weight [WW])
- *b*_*t*_ is the contemporary group effect
- *u*_*t*_ is the calf genetic effect (fit only for weaning weight) and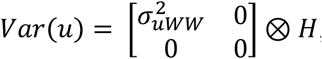, where 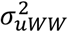 represents the genetic variance for the direct effect of weaning weight
- *m*_*t*_ is the cow genetic effect and 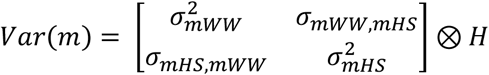, where 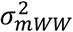 represents the genetic variance for the maternal effect of weaning weight and 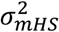 genetic variance for hair shedding
- *mpe*_*t*_ is the cow permanent environment effect and 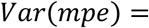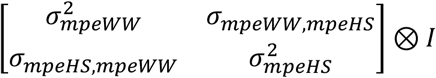 represents the, where 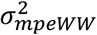 represents the permanent environmental variance for the maternal effect of weaning weight and 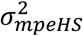 represents the permanent environmental variance for hair shedding
- *e*_*t*_ is the random residual and 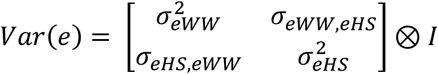
- *X*, *Z*_1_, *Z*_2_, and *Z*_3_ are incidence matrices relating the elements of *Y* to *b*, *u*, *m*, and *mpe*, respectively

We also evaluated the phenotypic relationship between dam hair shedding score and calf weaning weight using data from a subset of 6,448 dams with both hair shedding scores and calf weights recorded in at least one year (*n* = 9,092 score/weight pairs) by calculating the estimated change in calf weaning weight as a function of dam hair shedding score using four separate simple linear regression models. In the first two models, unadjusted calf weaning weight was regressed on unadjusted dam hair shedding score. Using weaning weight unadjusted for age in days captures increased gain from an earlier birth date (older when weighed), which might be an indicator of increased fertility for earlier shedding cows. In the other two models, 205-day, age-of-dam, and contemporary group solution adjusted calf weaning weight was regressed on un-adjusted dam hair shedding score. Both the unadjusted weaning weight and adjusted weaning models were fit separately for all available data, dams explicitly recorded as grazing toxic fescue, and dams explicitly recorded as not grazing toxic fescue.

### Genome-wide association

In order to evaluate the genetic architecture of hair shedding and identify variants contributing to hair shedding score breeding values, we performed a single-SNP genome-wide association analysis using SNP1101 v.1 (30). The breeding values calculated above using AIREMLF90 were de-regressed and used as the phenotype such that each of the 3,783 animals had one record. The de-regressed breeding values were weighted by their reliability 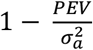, where 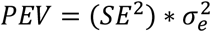 and 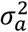 and 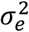 are the estimated additive genetic and residual variances for hair shedding score, respectively. Heritability was constrained to 0.40 and the genomic relatedness matrix used to control for family structure was calculated using the VanRaden method (31).

- Using the UMD 3.1 bovine genome assembly (32) coordinates and annotations, we searched genes within 50 Kb of SNP variants with a genome-wide *q*-value lower than 0.05. The size of our search space was determined based on the density of our marker set, and the resulting gene list was used as input for cluster enrichment analysis within ClueGO v.2.5.6 (33). KEGG pathways and biological process gene ontologies with at least 4 associated genes were considered for search terms. We also searched for protein-protein interaction between genes in our list using STRING v.10 (34), considering co-expression, experimental data, and curated databases as active interaction sources.

## Results

### Contemporary group definition

The results of the age-in-years model suggest a non-linear effect of age with larger effect sizes in two-year-olds, three-year-olds, yearlings, and old cows relative to mature cows (Figure 3a). Both the BIF age classes model and the four age classes model had lower AIC values than the null model with no age effect (38912.38, 38906.17, and 38983.31 respectively). Likelihood ratio test results indicate a better fit of the four class model over the null (*−log10(p)* = 8.899) and no improvement in model fit using BIF age classes over four age classes (*−log10(p)* = 0). Therefore, we chose to classify age using the simpler four age class model in all downstream analyses in order to maximize contemporary group size.

**Figure 3.**
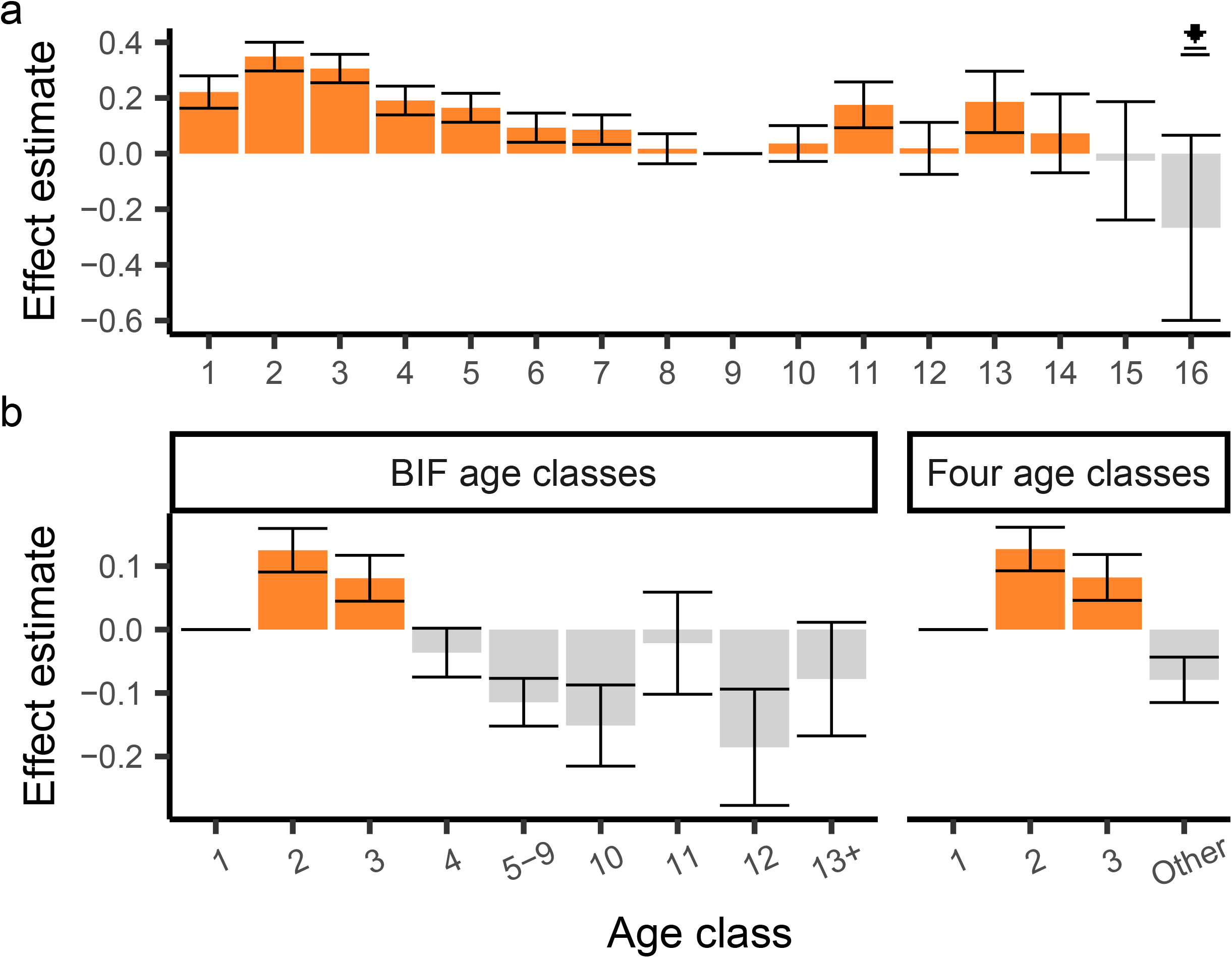
Estimates of the effect of age on hair shedding score. **a.** The effect of age in years on hair shedding score appears to be non-linear, following a U-shaped pattern. **b.** Comparison of effect estimates using BIF age-of-dam classifications or four age classes. Error bars represent standard error. Age groups with at least five observations are plotted.

When treated as separate traits, hair shedding while grazing and not grazing toxic fescue had similar heritability estimates (Table 1) and a high genetic correlation (*r*_*g*_= 0.93). Further, the Pearson correlation between breeding values grazing and not grazing toxic fescue was 0.99.

**Table 1.**
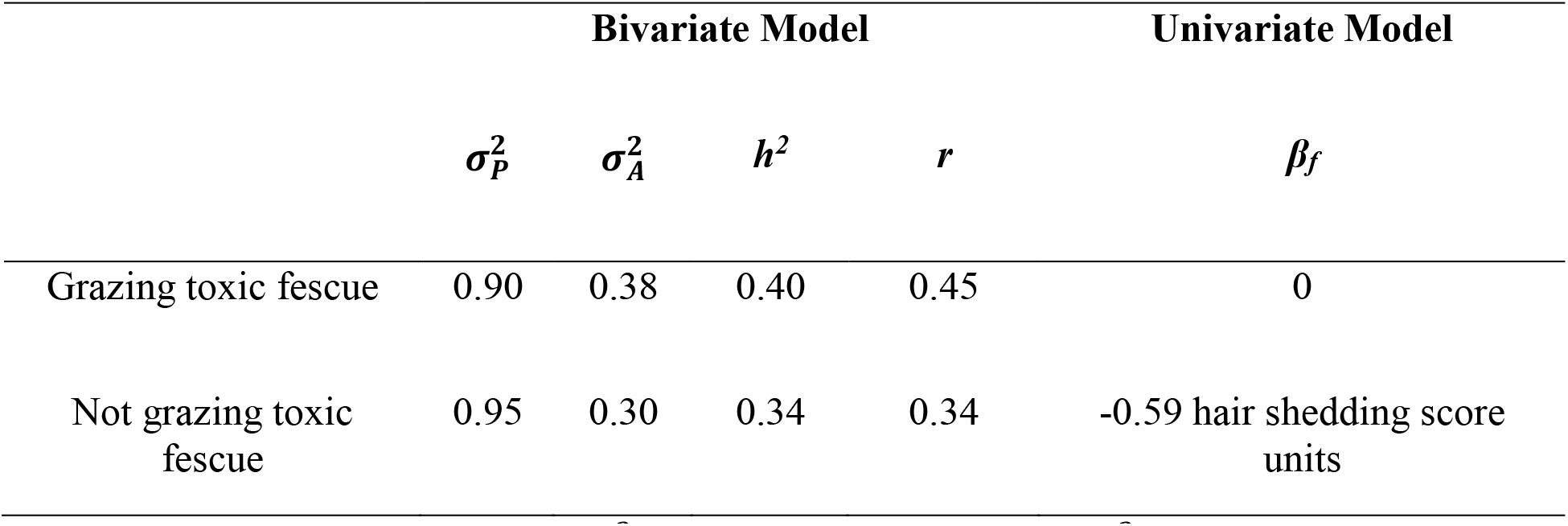
Comparison of genetic parameters estimated using cattle grazing and not grazing toxic fescue. The estimated phenotypic variance 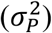, additive genetic variance 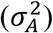, narrow sense heritability (*h*^2^), and repeatability (*r*) from a bivariate model and fixed effect of grazing versus not grazing fescue from a univariate model. Additive genetic variance, heritability, and repeatability are higher for hair shedding recorded while grazing toxic fescue when treated as a different trait from hair shedding while not grazing toxic fescue. When fescue grazing status is fit as a fixed effect in a univariate model, the estimated effect of toxic fescue on hair shedding score (*β*_*f*_) is also higher (i.e. later shedding animals).

|  | Bivariate Model |  |  |  | Univariate Model |
| --- | --- | --- | --- | --- | --- |
| | $\sigma_P^2$ | $\sigma_A^2$ | $h^2$ | $r$ | $\beta_f$ |
| Grazing toxic fescue | 0.90 | 0.38 | 0.40 | 0.45 | 0 |
| Not grazing toxic fescue | 0.95 | 0.30 | 0.34 | 0.34 | -0.59 hair shedding score units |

The total phenotypic variation in hair shedding grazing toxic fescue was slightly higher than hair shedding not grazing toxic fescue, suggesting reduced peripheral blood flow caused by fescue toxicosis is more detrimental to hair shedding than heat stress alone (Table 1). The fixed-effect model solutions support this conclusion (*β*_*f*_ = 0 vs. −0.59 hair shedding score units for grazing and not grazing toxic fescue, respectively). Further, the estimated permanent environment effect (and therefore estimated repeatability,*r*) was much higher for hair shedding while grazing toxic fescue (Table 1).

### Genetic parameters and prediction

Using all available data, the estimated narrow-sense heritability 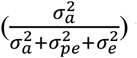 was 0.40, and the estimated repeatability 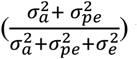 was 0.44. These estimates are similar to those previously found in Angus cattle based on pedigree relatedness (21).

Across ten iterations, 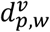 averaged 0.25, ranging from 0 to 1.48. In the absence of bias introduced by incorrect estimation of the genetic trend, this value is expected to be zero. Estimates of 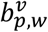 ranged from to 1.05, suggesting minimal dispersion of breeding values (Figure 4). Prediction accuracy 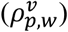 ranged from 0.70 to 0.73.

**Figure 4.**
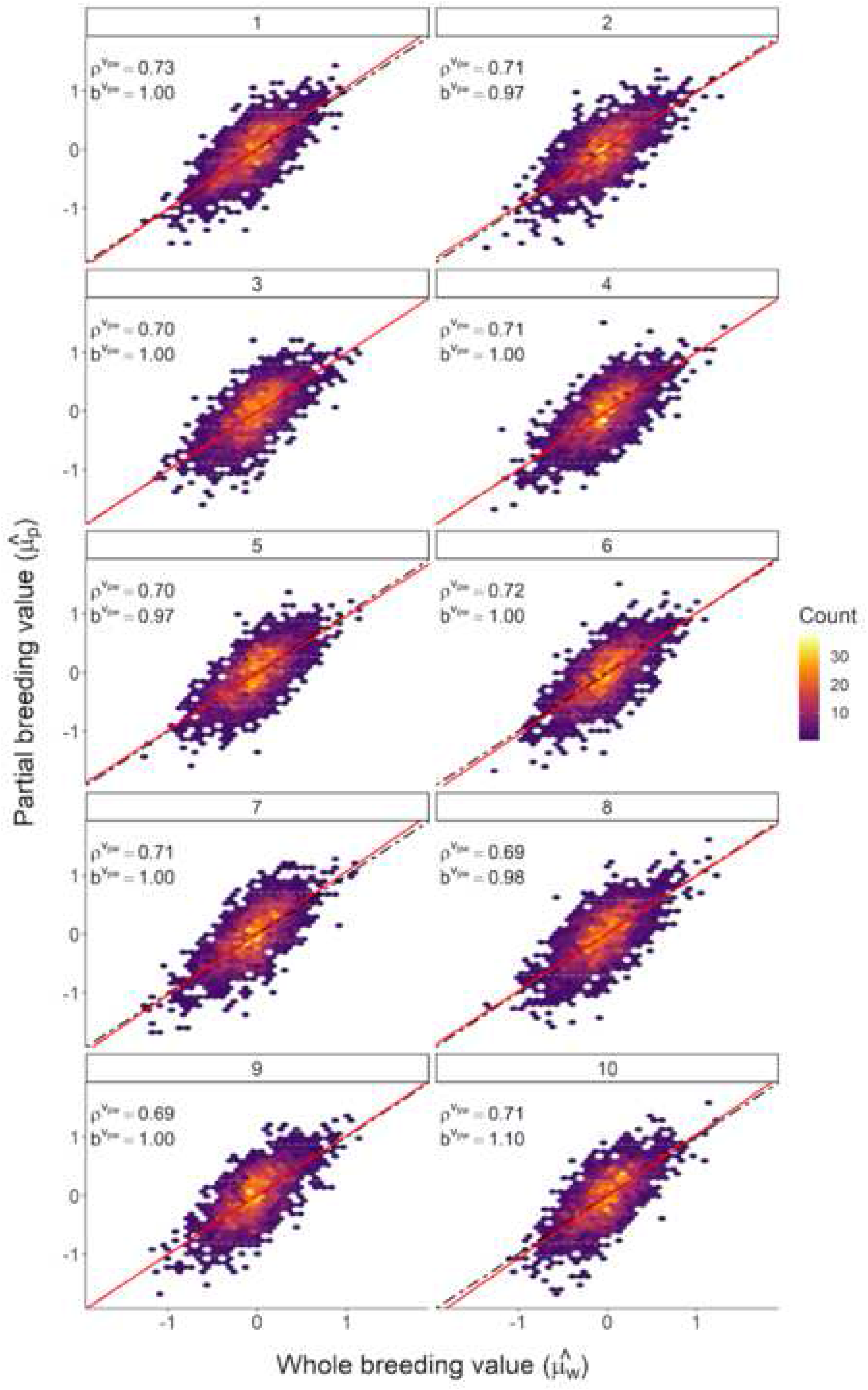
Linear regression evaluation of breeding values. Comparison of breeding values estimated using all available data 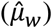 and breeding values estimated using a reduced dataset 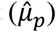 across ten iterations within validation animals. The solid red line represents 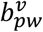 and the dotted black line represents the expectation of 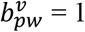 in the absence of dispersion.

### Weaning weight

The effects of heat stress on pre-weaning growth are well characterized in beef and dairy cattle. Heat stress impacts calf performance most severely via reduced milk production in the dam (35). Fescue toxicosis induces reduced milk production in a similar fashion (36). Therefore, we quantified the phenotypic and genetic correlations between hair shedding score and weaning weight.

All three bivariate models suggest a moderately negative genetic correlation between weaning weight and hair shedding score. In the model using all available data, the estimated *r*_*g*_ between the maternal component of weaning weight and hair shedding was −0.19 (Table 2). When the data were stratified by dam toxic fescue grazing status, this estimate increased in magnitude slightly for both grazing and not grazing toxic fescue (*r*_*g*_ = −0.25 and −0.28, respectively). For dams not grazing toxic fescue, the *r*_*g*_ between the direct and maternal effect of weaning weight fell near commonly reported estimates (*r*_*g*_ = −0.29; (37)) but was much higher for dams grazing toxic fescue (*r*_*g*_ = −0.63) and for all possible dams (*r*_*g*_ = −0.43) (Table 2). The *r*_*g*_ between the direct effect of weaning weight and hair shedding varied from −0.10 (dams not grazing toxic fescue) to −0.03 (all possible data) to zero (dams grazing toxic fescue).

**Table 2.**
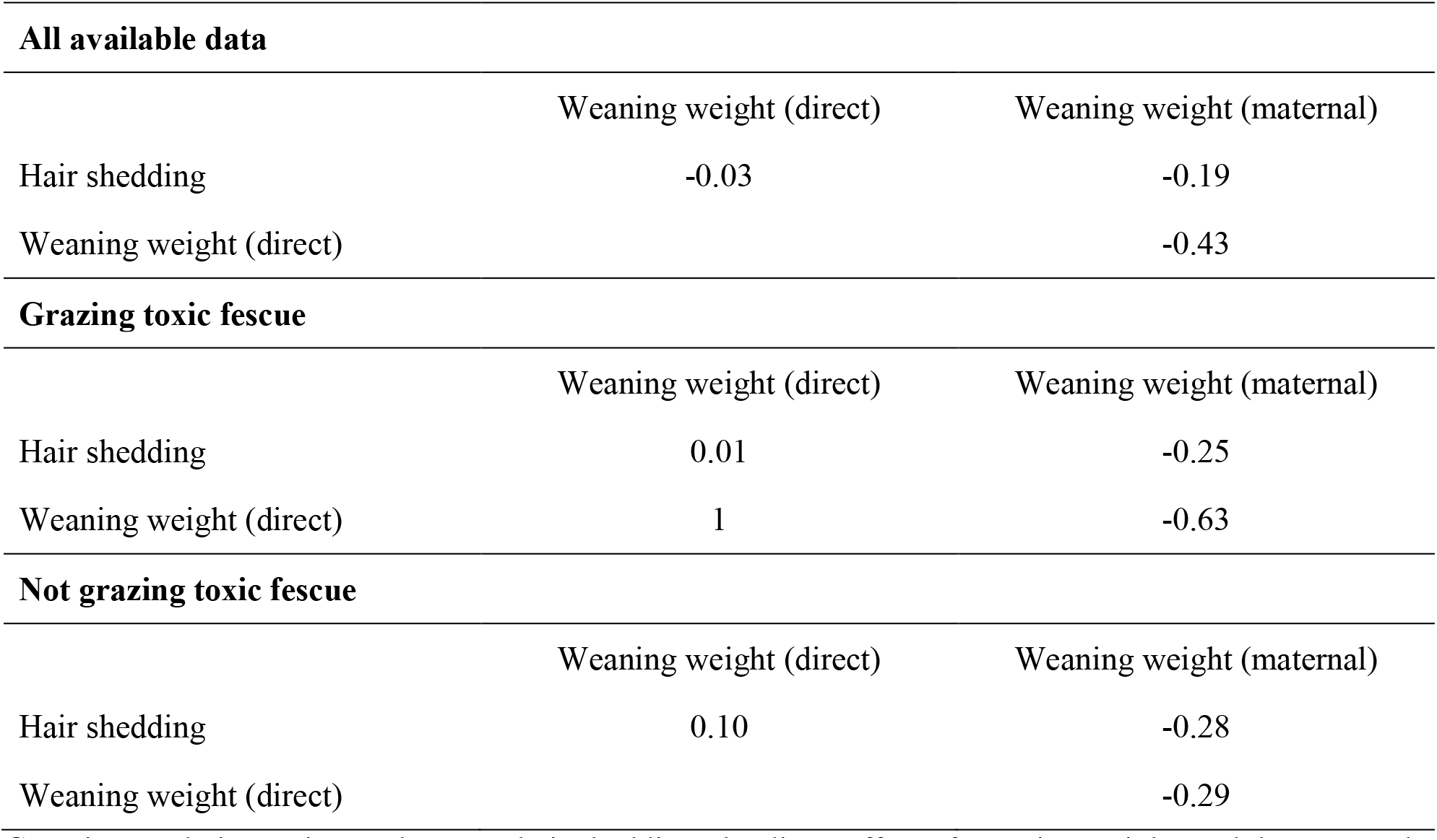
Estimated genetic correlations between dam hair shedding and calf weaning weight. Genetic correlation estimates between hair shedding, the direct effect of weaning weight, and the maternal effect of weaning weight vary across toxic fescue grazing statuses.

| **All available data** |  |  |
| --- | --- | --- |
|  | Weaning weight (direct) | Weaning weight (maternal) |
| Hair shedding | -0.03 | -0.19 |
| Weaning weight (direct) |  | -0.43 |
| **Grazing toxic fescue** |  |  |
|  | Weaning weight (direct) | Weaning weight (maternal) |
| Hair shedding | 0.01 | -0.25 |
| Weaning weight (direct) | 1 | -0.63 |
| **Not grazing toxic fescue** |  |  |
|  | Weaning weight (direct) | Weaning weight (maternal) |
| Hair shedding | 0.10 | -0.28 |
| Weaning weight (direct) |  | -0.29 |

In the simple linear models predicting unadjusted weaning weight from dam hair shedding score, unadjusted calf weaning weight was estimated to decrease by 1.30 kg with every unit increase in hair shedding score using all available data, by 3.22 kg for dams grazing toxic fescue and by 5.08 kg for dams not grazing toxic fescue. Slope estimates from the simple linear models predicting adjusted weaning weight from dam hair shedding score were more modest but also negative. Adjusted calf weaning weight was estimated to decrease by 1.45 kg using all possible data, by 2.47 kg among dams grazing toxic fescue and by 1.11 kg among dams not grazing toxic fescue with every unit increase in hair shedding score (Figure 5).

**Figure 5.**
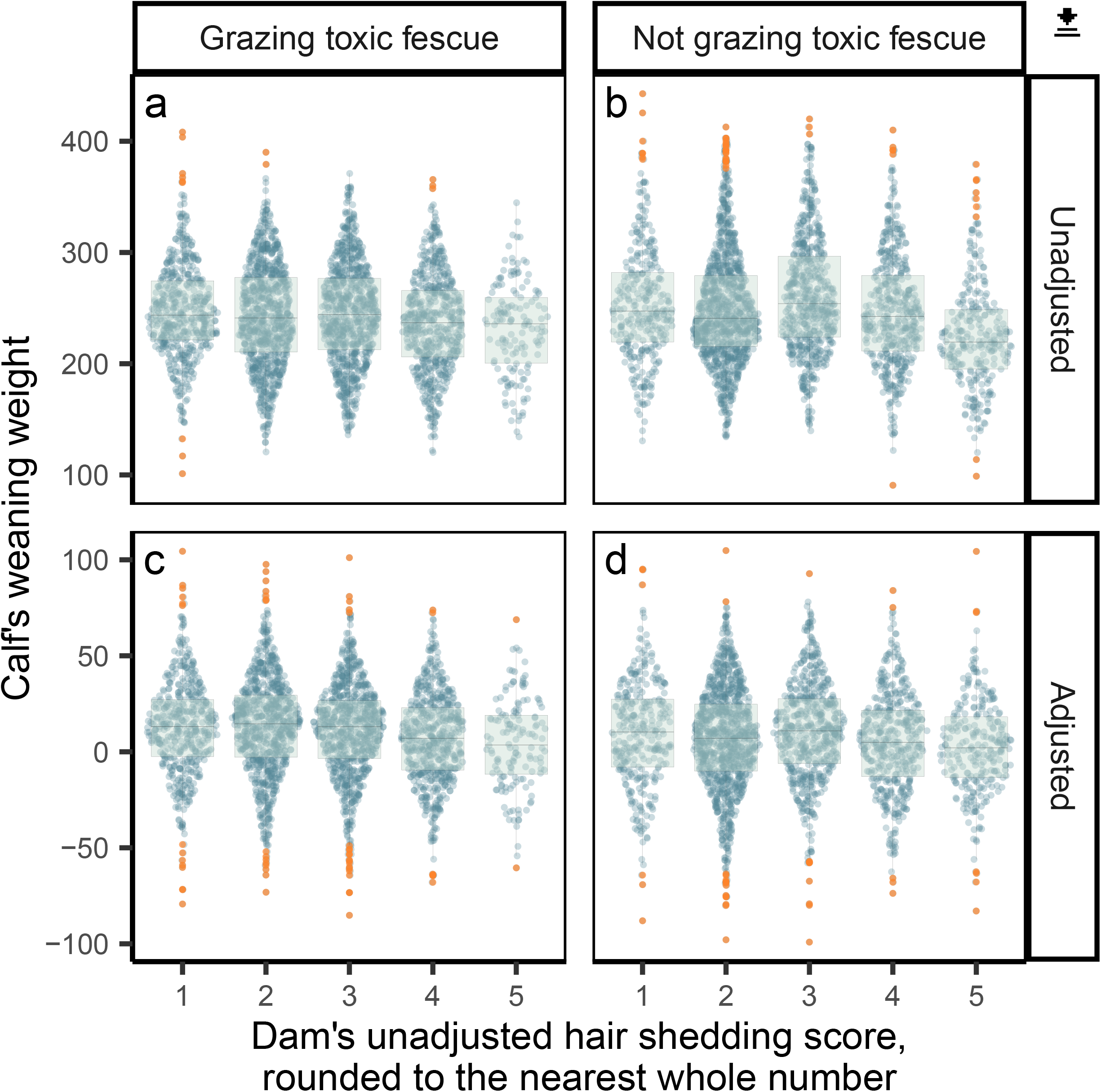
Comparison of dam’s hair shedding score to the weaning weight of her calf. The effect of a dam’s hair shedding score on the unadjusted (**a, b**) and adjusted **(c, d**) weaning weight of her calf with outlier weaning weights highlighted in orange. Regardless of fescue grazing status, there is very little difference in calf weaning weight between dams with hair shedding scores 1, 2, and 3.

### Genome-wide association analysis

We found 176 variants passing a genome-wide false discovery rate threshold of 0.05 and 56 variants passing a false discovery rate threshold of 0.01 (Figure 6). Of these 176 variants, 33% reside on BTA5. Two hundred and six unique genes were found to be within 50 Kb of passing variants. The two strongest associations were observed within *CEP290*. Perhaps interestingly, we identify several members of the KRT gene family (*KRT1, KRT3, KRT4, KRT76, KRT77, KRT78*, and *KRT79*), which are involved in creating structural epithelial cells like hair, near our largest peak.

**Figure 6.**
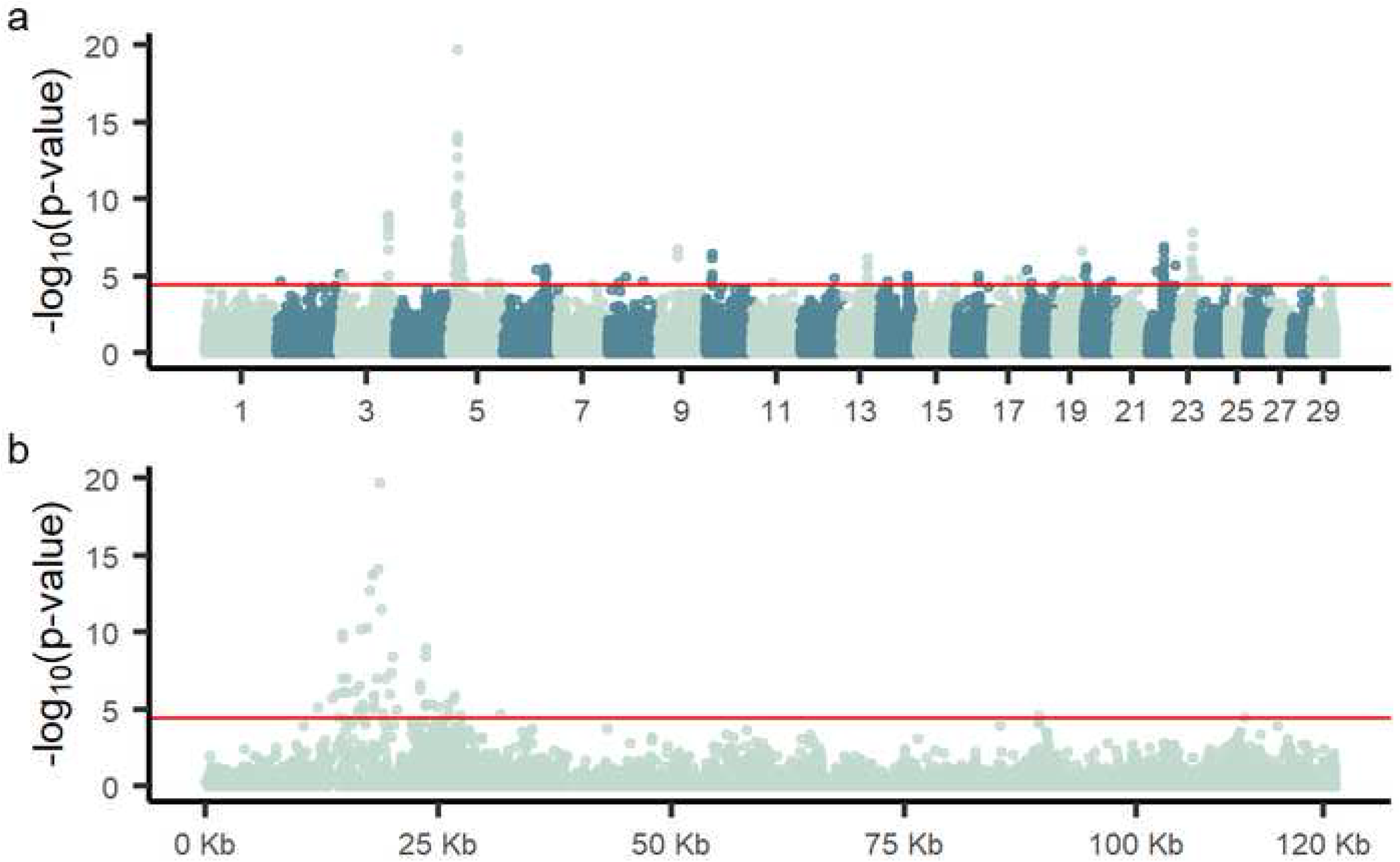
Manhattan plot of variants associated with hair shedding. Using de-regressed hair shedding score breeding values in SNP1101 single-SNP regression, we find 176 variants significantly associated with hair shedding (FDR < 0.05, red line) (**a**). Of these 176 variants, 33% reside in a peak on BTA5 (**b**).

We found significant enrichment (Benjamini-Hochberg corrected p-value < 0.05) for pathways involved in virus-host interaction, fat cell differentiation, prolactin signaling, cellular response to starvation, Vasopressin-regulated water reabsorption, and other biological processes (Table 3). We also found more protein-protein interactions than expected (PPI enrichment p-value = 0.00462) and enrichment for PFAM protein domains “keratin type II head” (FDR = 8.89e-06), “somatotropin hormone family” (FDR = 8.09e-05), and “intermediate filament protein” (FDR = 0.00064).

**Table 3.**
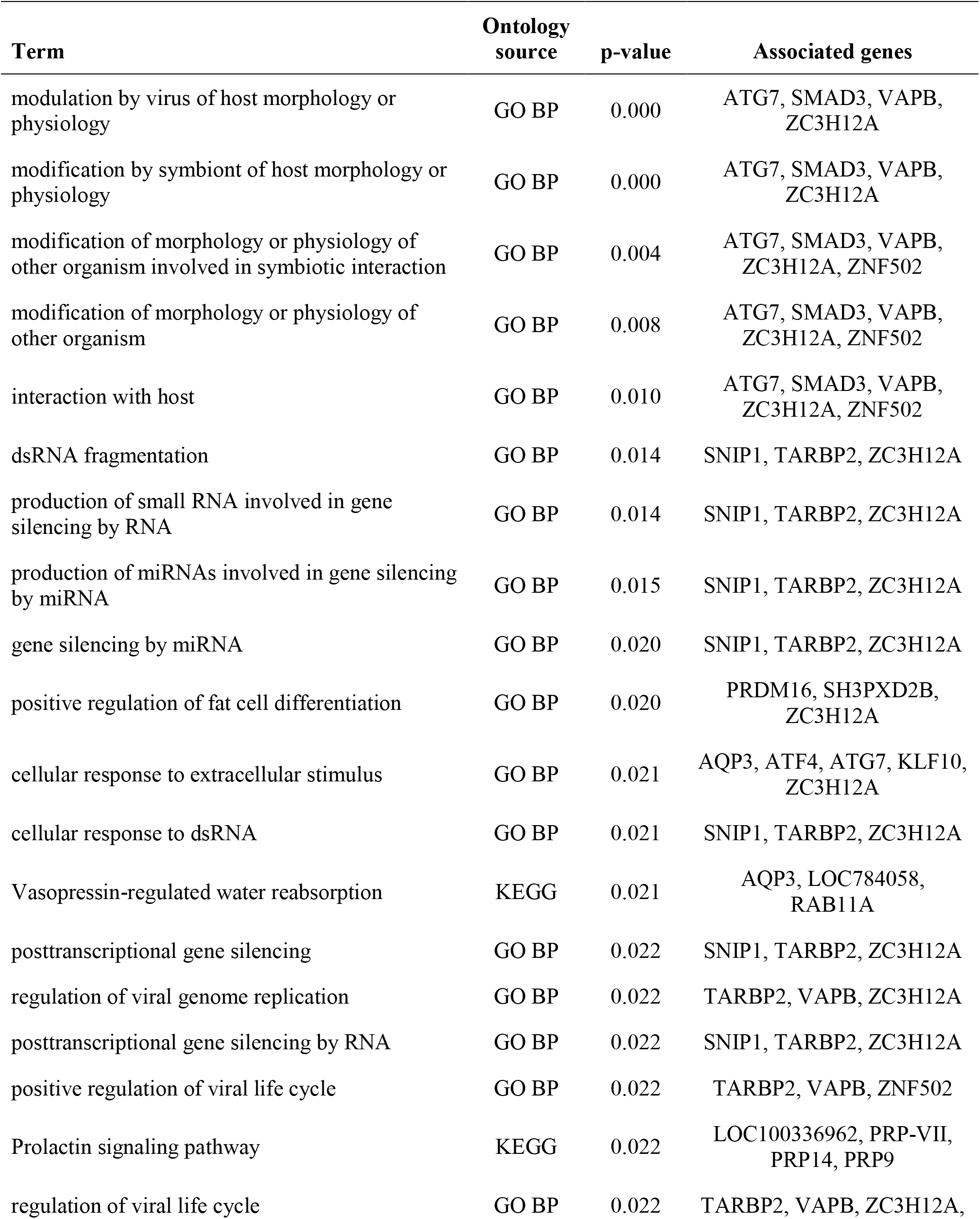
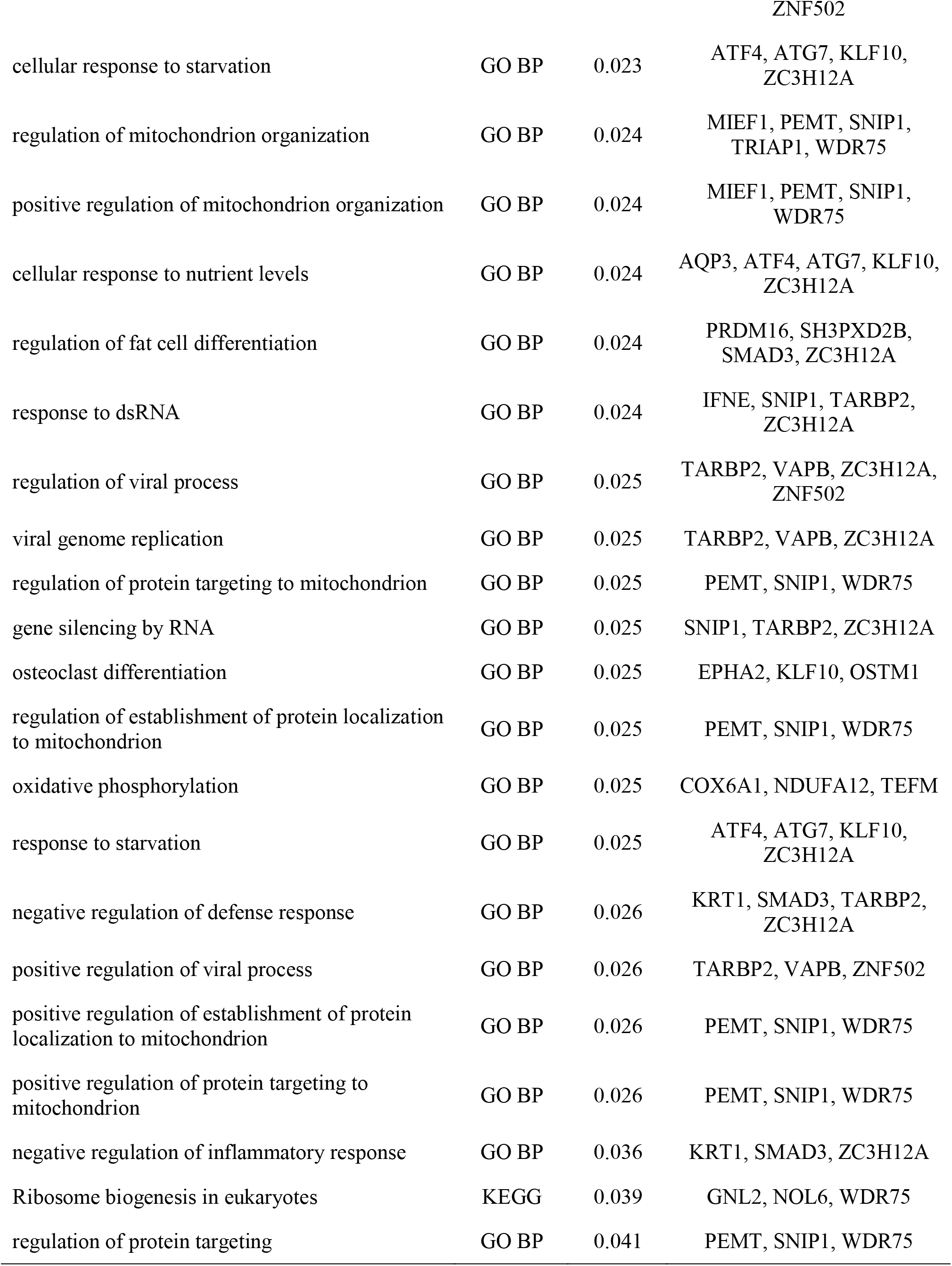
Terms significantly associated with genes within 50 Kb of hair shedding GWAA variants with FDR < 0.05. We find enrichment for pathways involved in virus-host interaction, response to starvation, prolactin signalling, and other biological processes. P-value are corrected for multiple testing using Benjamini-Hochberg methodology. Enrichments represent gene ontology biological process (GO BP) or Kyoto Encyclopedia of Genes and Genomes pathways (KEGG).

## Discussion

The expression of a phenotype is not always consistent across lifespan (38,39). We found that the relationship between age and hair shedding is non-linear with young cows, especially 2-year-olds and 3-year-olds, displaying higher hair shedding scores than their older herd mates. This is in line with expectations, as young cows require increased energy expenditure associated with continued growth (40) and the new stress of lactation (41). To a lesser extent than young animals, cows 10 to 13 years old tended to have higher hair shedding scores. A similar U-shaped relationship between age and molt date has been observed in other ungulate species (8) and was reflected in the effect size estimates from the BIF age class model (Figure 2b). Cows are typically culled from the herd or die after 10 to 11 years of age (42,43). Thus, effect estimates for cows older than 12 reflect a selected sample. However, the early shedding estimates for these very old cows support early hair shedding as an important characteristic of longevity, especially in heat-stressed environments.

Though our results suggest a high correlation between hair shedding score breeding value across toxic fescue grazing status, we estimate a slightly higher heritability and much larger effect of permanent environment among cattle grazing toxic fescue than those not grazing toxic fescue. Stress can increase the potential for expression of genetic differences (38), which could result in the higher heritability observed among cattle grazing toxic fescue. The disparity found in permanent environment estimates might be explained by a higher contribution of non-additive genetic effects (i.e., epistatic and dominance effects) to variation in hair shedding while grazing toxic fescue versus while not grazing toxic fescue. It’s also possible that certain permanent environmental effects (i.e., physiological differences between the ability of animals to shed their winter hair) are manifest when cattle graze infected tall fescue. Most likely, the increased permanent environment effect estimate is a reflection of the accrual of physiological damage from long-term fescue toxicosis. The medial layer of blood vessels tends to be thickened in animals suffering from fescue toxicosis, which (44) linked to hyperplasia of the smooth muscle. Repeated exposure to ergovaline also increases venous contractile response, suggesting bioaccumulation (45).

Typically, measurements of the same trait across different environments that result in genetic correlations below *r*_*g*_ = 0.70 are considered “very different” (46). Hair shedding scores recorded while grazing toxic fescue versus while not grazing toxic fescue have an *r*_*g*_ of 0.93, suggesting minimal re-ranking of breeding values. However, the magnitude of difference in permanent environment effects found here may justify treating hair shedding grazing and not grazing toxic fescue as separate traits in research studies examining physiological or non-additive genetic effects. For the implementation of the American Angus NCE, we’ve chosen to minimize the effect of toxic grazing status by including it in the definition of contemporary groups. Many biotic and abiotic factors affect the prevalence of toxicity-inducing ergot alkaloids within forage, including moisture, reproductive status, soil nitrogen, and most notably, temperature (47). Previous work suggests that animals must ingest a threshold level of ergot alkaloids before fescue toxicosis symptoms become evident (48). However, in these analyses, toxic fescue grazing was treated as a binary producer-reported status in the absence of quantitative measures of ergot alkaloid levels, which may affect the interpretation of our results. Further, we did not account for the effect of grazing toxic fescue in previous years.

Our enrichment results identified pathways associated with prolactin signaling, which is a well-known modulator of seasonal hair shedding and hair growth as well as milk production (4). Further, low serum prolactin level is often used as an indicator of fescue toxicosis (49). Gray et al. (2011) suggested that the negative relationship they found between calf weaning weight and dam hair shedding was due in part to differences in serum prolactin level. Our results support that conclusion. While the genetic correlation found here using all possible data between a dam’s hair shedding score and the weaning weight of her calf is moderate, it’s nearly three times less than the previous estimate reported by Gray et al. (2011) ($r_g$$ = - 0.58), which was identical to the correlation reported by Turner & Schleger (1960) for a calf’s own hair shedding score and its post-weaning gain. This is likely due in part to our use of a slightly different phenotype. Turner & Schleger (1960) used an expanded 7-point scoring system, while Gray et al. (2011) used the same scoring system but categorized dams based on the month of the year that they first achieved a hair shedding score of 3 (about 50% shed; Figure 1c). We also considered the relationship between hair shedding score and the maternal effect of weaning weight rather than the direct effect of weaning weight.

Another possibility could be confounding environmental effects. The relationship between dam hair shedding score and calf weaning weight was also different across toxic fescue grazing statuses, and when toxic fescue grazing statuses are modeled separately the *r*_*g*_ between hair shedding score and the maternal effect of weaning weight increases relative to the *r*_*g*_ estimated using all data. This is similar to results found by Hoff et. al (2019), where the accuracy of bovine respiratory disease (BRD) genomic prediction was higher when data were analyzed stratified by state than when data were analyzed all together. The authors postulated that the discrepancy in prediction accuracy was likely due to the prevalence of different BRD-causing pathogens between environments (50). Similarly, our results suggest that the relationship between hair shedding and other production traits may be environment- or context-specific.

In the four phenotypic regressions of calf weaning weight on dam hair shedding score, dam hair shedding while grazing toxic fescue was estimated to have the largest effect on adjusted weaning weight, but not unadjusted weight. When contemporary grouping is fit as a fixed effect in BLUP, the resulting contemporary group solution can be interpreted as a metric of environmental stress (51,52). Larger contemporary group solutions indicate a greater advantage to the phenotype from the environment, including plane of nutrition and management practices. The disparity we find between adjusted and unadjusted weaning weight results can be explained by smaller contemporary group solutions among calves whose dams grazed toxic fescue. Indeed, we find the mean contemporary group solution among calves whose dams did not graze toxic fescue to be 20.75 kg higher than that of those whose dams did graze toxic fescue (258.95 and 238.20 kg, respectively).

The negative genetic correlation often found between the maternal and direct genetic effects of weaning weight has puzzled researchers since the first large scale national cattle evaluations, with some suggesting that it is artifactual and others that it is a reflection of real biological phenomena (53). We found that the magnitude of this genetic correlation varied across toxic fescue grazing statuses, with dams grazing toxic fescue showing a more negative correlation (−0.63) than dams not grazing toxic fescue (−0.29) (Table 2). There are several potential explanations for this result. First, the variation we find in genetic correlations between maternal and direct weaning weight could be a result of sire-by-herd and sire-by year interactions (54). These interactions can arise via multiple avenues, including genotype-by-environment interactions, selective data reporting, and preferential management of the progeny of certain sires. If this interaction were larger in certain herds, our estimates would be skewed. Alternatively, it’s possible that our results reflect the effect of fescue toxicosis on dam nutrient partitioning. Our enrichment analysis identified multiple pathways involved in response to nutrient levels, response to starvation, and fat cell differentiation, which could support this conclusion. During the initiation of lactation, mammals draw upon their own energy reserves in order to meet increased metabolic demand (55,56), which implies genetic antagonism between maternal and direct weaning weight (37,53). The nutrient partitioning process is influenced by stress. For example, Rhoads et al. (2008) (57) demonstrated that decreased feed intake explains only part of the reduction in milk yield found in heat-stressed dairy cows, indicating further changes in metabolism and partitioning of nutrients in response to hyperthermia.

Although associations with a FDR less than 0.05 were found on 20 of 29 autosomes and associations with an FDR less than 0.01 were found on seven chromosomes, one third of the associated variants resided on BTA5. Among these, ten variants were found near or within members of the keratin gene family.

Particularly, *KRT1, KRT3, KRT4, KRT77, KRT78*, and *KRT79* form a protein-protein interaction network, the orthologs of which are co-expressed in other species during the formation of intermediate filament proteins. However, it’s possible that significant variants near and within keratin genes are simply an artifact of extensive linkage disequilibrium (Figure 6b). Using the current sample, this result is difficult to disentangle.

The two most significant associations were both contained in *CEP290*. In humans, mutations in *CEP290* cause abnormal photoreceptors (58,59). Photoreceptors affect an animal’s ability to detect changes in seasons (60), and changes in photoreceptors could have large impacts on this function. Mutations in *CEP290* affect cilia formation, and are believed to interact with Bardet–Biedl syndrome (BBS) proteins (61).

Recently, *BBS1* was associated with local adaptation in Red Angus cattle (18). Further, the strength of these associations on chromosome 5 could be due to multiple causal mutations (62) in this locus from 12 Mb to 28 Mb.

## Conclusions

We developed a prototype genetic evaluation for early-summer hair shedding in American Angus cattle in order to enable genetic selection for heat tolerance. In agreement with previous research (20,21), we find early summer hair shedding to be moderately heritable. We also identified variants associated with biological pathways like prolactin signaling, response to starvation, and keratin formation that contribute to genetic variation for hair shedding score.

Weaning weight and hair shedding score appear to be negatively correlated, and we find evidence for a greater impact of hair shedding score on performance for cows experiencing heat stress alone compared to cows grazing toxic fescue. Further investigation of the relationship between hair shedding and other symptoms of fescue toxicosis (like reduced fertility) as well as the functional biology of hair shedding is necessary in order to determine the appropriateness of using hair shedding scores as an indicator trait for tolerance to fescue toxicosis. However, many of our results support the use of hair shedding scoring as a barometer of cow health in addition to other routinely collected phenotypes like body condition score.

### List of abbreviations

- AAA = American Angus Association
- BIF = Beef Improvement Federation
- NCE = National Cattle Evaluation
- WW = weaning weight
- HS = hair shedding

## Declarations

(please see our editorial policies for more information: https://www.biomedcentral.com/getpublished/editorial-policies#ethics+and+consent). If any of the sections are not relevant to your manuscript, please include the heading and write ‘Not applicable‘ for that section.

### Ethics approval and consent to participate

Because phenotypic records and tissue samples were collected as part of routine livestock production practices, an ACUC exemption was granted (University of Missouri ACUC Protocol Number EX 8596).

### Consent for publication

Not applicable.

### Availability of data and materials

Datasets supporting the conclusions of this article are available for non-commercial use via a data use agreement (DUA) with the American Angus Association.

### Authors’ contributions

JED and HJD conceptualized and designed the research. HJD and HYI managed data acquisition, storage, and retrieval. DL performed imputation. HJD performed variance components, genetic prediction and association analyses. HJD, SPM, and JED interpreted results. HJD, SPM, and JED wrote the initial version of the manuscript, which was edited by all authors.

## Acknowledgements

We acknowledge the many farmers and ranchers who collected data as part of this project. We also acknowledge the data collected by research teams of Trent Smith at Mississippi State University and Joe Cassady at North Carolina State University who contributed data to Angus Genetics Inc. We appreciate assistance from Daniela Lourenco and Kelli Retallick in using the BLUPF90 suite of programs and for discussion of analyses. Assistance from Mike MacNeil, Jason Kenyon, Lou Ann Adams, and Cody Combs in accessing and organizing AAA data was also greatly appreciated.

